# Real time evaluation of the liver microcirculation by whole organ machine perfusion within an MRI system

**DOI:** 10.1101/2025.05.08.652602

**Authors:** Zainab L Rai, Natalie A Holroyd, Morenike Magbagbeola, Emre Doganay, Katie Doyle, Lucy Caselton, Agositino Stili, Danail Stoyanov, Simon Walker-Samuel, Brian R Davidson

## Abstract

**Objectives:** Machine perfusion of organs outside of the body is a growing area of research with significant applications in the fields of organ preservation and transplantation, but more widely it offers a new approach to study disease processes and to evaluate new therapeutics and devices. Magnetic Resonance Imaging (MRI) allows for non-invasive assessment of organ structure and function, enabling quantitative measurement of tissue perfusion and microstructure. In this study, we demonstrate that MR imaging sequences can be obtained from machine-perfused porcine livers using a modified perfusion rig for MR compatibility and highlight the quantitative measures that can be obtained through this methodology.

**Materials and Methods:** 7 porcine livers were retrieved fresh from the abattoir using a previously published protocol and following transport in cold preservative underwent perfusion with oxygenated autologous blood inside a 3T clinical MRI scanner using a custom modified perfusion rig. Multiple MR imaging sequences were acquired: T2-weighted imaging, Diffusion Weighted Imaging and Dynamic Contrast Enhanced imaging following injection of Gadolinium dye into the portal vein and hepatic artery. Histological analysis was performed to assess preservation injury to the liver. Control samples for histology were obtained from livers with similar preservation periods but preserved in standard cold storage on ice (Static Cold Storage).

**Results:** Concurrent MR imaging and machine perfusion were successfully performed, allowing dynamic measurement of tissue perfusion to be obtained in ex vivo livers, including calculation of gadolinium contrast enhancement curves and Apparent Diffusion Coefficient maps. Segmentation of vessels down to a radius of 0.45mm allowed detailed morphological analysis of the vascular network, including extraction of clinically relevant parameters such as vessel tortuosity. Histological evaluation showed better preservation of the hepatic acinar structure in perfused than non-perfused livers.

**Conclusions:** Our results demonstrate that MR imaging of machine-perfused organs enables high-resolution quantitative evaluation of whole-organ vascular morphology and flow dynamics. This platform provides opportunities to study vascular pathology in diseased human organs and evaluate novel therapeutic interventions, with particular relevance for drug-delivery strategies.

## Introduction

Machine perfusion of whole organs has been developed as a means of preserving human organs for transplant^1–4^. Increasingly, perfused organs are also being explored for research purposes, enabling researchers to better understand human organ physiology and pathophysiology^2,5,6^. In this setting, machine perfused organs offer a unique, human model system for the development, evaluation and safety testing of new devices and drugs for a spectrum of acute and chronic diseases^7–9^. Previous studies have reported on a custom built novel sub-normothermic machine perfusion system with inline sensors, developed for the purpose of creating a perfused whole organ research model^10,11^.

Magnetic Resonance Imaging (MRI) is widely used clinically to non-invasively assess the liver, offering quantitative measurements of perfusion and tissue microstructure^12^. In this study, we demonstrate the use of MRI to provide comprehensive, quantitative evaluation of vascular structure and function in machine-perfused porcine livers. Using a previously described sub-normothermic liver perfusion circuit^10^, modified to allow perfusion within an MRI scanner, we evaluate the utility of ex-vivo perfused liver MRI as a research platform by (1) reconstructing vessel networks to extract morphological characteristics, (2) quantifying microstructural properties using Diffusion Weighted Imaging (DWI), and (3) measuring flow within the vasculature using Dynamic Contrast-Enhanced (DCE) MRI following injection of a contrast agent. Additionally, we confirm organs were preserved throughout the perfusion period by conducting histological analysis and comparing to organs kept for a similar period under Static Cold Storage (SCS).

DWI and DCE-MRI imaging sequences were chosen for this study as both can be used in patients to assess diverse liver pathologies, such as characterising a focal liver lesion^13,14^ and grading of liver fibrosis^15,16^. In DWI, the MRI scanner is sensitised to the diffusive motion of water in tissue, and by mathematically modelling the resultant signal, parameters linked to microstructure (e.g. cellularity and vascularisation) can be estimated. Likewise, perfusion MRI is a key clinical modality for determining the nature of focal liver lesions, the success of local tumour ablation and in the development of anti-angiogenic agents for cancer treatment^17–19^. Dynamic contrast-enhanced MRI (DCE-MRI) is used to assess perfusion of organs by injecting a paramagnetic contrast agent and imaging the subsequent signal enhancement. Model-based or model-free methods are utilised to analyse the contrast concentration-time curve in liver lesions or parenchyma^20,21^.

### Imaging of machine perfused organs

Existing work in the field of biomedical imaging of perfused organs has mainly focused on assessing organ viability and suitability for transplant, for example using DWI to detect warm ischaemic damage in machine-perfused kidneys^22^. Similarly, Arterial Spin Labelling (ASL) MRI has been used to assess perfusion quality in porcine and human kidneys^23,24^. Hyperpolarised MR, using [1-^13^C]pyruvate contrast, has been used to evaluate metabolic function in porcine hearts to assess graft quality^25^. In liver, contrast-enhanced ultrasound has been employed to detect reperfusion defects in porcine organs^26^. Beyond assessing organs for transplant, some work has been done to develop machine-perfused organs as experimental models. For example, Pelgrim et al. validated measures of myocardial ischemia and blood flow determined using dynamic Computed Tomography in machine perfused porcine hearts^27^. Likewise, an MR compatible perfused porcine heart model was developed by Vaillant et al. for investigation of haemodynamics and cardiac electrophysiology^28^. Notably, machine-perfused porcine livers have been used to study drug-drug interactions^29^; while this study did not involve biomedical imaging, it highlights the potential advantages of machine perfused organs as models for testing therapeutic agents.

This study builds on existing research by presenting a methodology for quantitative assessment of tissue perfusion and microstructure in ex-vivo perfused livers using MR imaging and highlighting the utility of this platform as a model for investigating human pathologies. To our knowledge, this is the first report of quantitative blood flow assessment in viable ex-vivo livers. ^22^

## Materials and Methods

### Development of mechanical perfusion system for use in the MRI magnet

The perfusion machine, designed as a research platform, has been previously described^10^. It was modified to allow organs to be perfused in the MRI magnet by housing the perfusion system in the insulated control room and connecting to the organ, which was housed in a water-tight plastic container in the MRI scanner, by silicone sealed tubing.

The perfusion system comprised of a reservoir with a capacity of holding up to 4 litres of autologous porcine blood, connected to a 2-liter pressurized canister containing medical-grade oxygen (BOC, Oxygen 1-E) to manually oxygenate the blood through a regulator (Figure1). A single peristaltic pump (Watson-Marlow, 520 DU) acted as a heat exchanger by supplying warm water through the oxygenator to achieve the desired temperature of the blood perfusing the organ. The reservoir was attached to an oxygenator, which was modified to allow it to be re-used on several occasions and to allow deep cleaning. A single centrifugal pump (PuraLev1200 MU) was employed to circulate the blood from the reservoir through the oxygenator, where it was warmed before perfusing both input vessels (hepatic artery and portal vein), the liver parenchyma, prior to returning to the reservoir via the hepatic veins. Along the input stream, between the reservoir and the liver, sensors were placed to monitor blood flow (Sonotec sonoglow, co.55/100), oxygen levels (PreSens, EOM-(t)-FOM), temperature (PreSens, Pt100), and pH (PreSens, EOM-(t)-FOM). Pressure sensors (PendoTech, Press-n-075) were positioned on both input streams (hepatic artery and portal vein), and a single pressure and flow sensor was placed on the output stream between the hepatic vein and the reservoir.

To achieve MRI compatibility, the reservoir, pumps, and sensors in the EM insulated MRI control room were connected via a 10-meter length of silicon tubing to the non-ferrous water-tight container, within which the input and output streams were connected to the major vessels of the organ. The tubing was primed using the peristaltic pump, to avoid air being pushed into the liver when the centrifugal pump was started.

For data acquisition and control, a Raspberry Pi Model 4 single-board computer (Raspberry Pi Ltd, UK) was integrated into the system. It facilitated real-time monitoring and recording of all sensor data through the MODBUS, RS-232, and RS-485 communication protocols. The software architecture was constructed on a Robotic Operating System (ROS2) framework, promoting modularity and scalability of the components.

### Perfusion Protocol

#### Organ retrieval, flushing, storage and transportation

Porcine livers and autologous blood were obtained from the abattoir using retrieval and transportation methods previously developed and reported^11^. The organs were retrieved and used in pairs. All organs were initially placed in a cooled and insulated storage box, maintained at a temperature of 4°C, for transportation. Upon arrival at the imaging facility, one organ from each pair was randomly allocated to remain in SCS for the duration of the experiment, thus acting as a control. Meanwhile, the other liver underwent perfusion and MRI imaging for a period of approximately 30 minutes, allowing direct comparison of tissue preservation injury in post-experimental histological evaluation.

#### Organ perfusion procedure in MRI system

The porcine livers undergoing perfusion were connected to the perfusion system as described previously^11^ then placed in a fluid-containing organ perfusion bowl within the MRI magnet. Autologous blood was collected and treated with heparin sodium (Pfizer) (4000 I/U per litre of blood) to avoid coagulation. Livers weighing less than 5kg were selected for the experiment, to ensure adequate perfusion with the volume of autologous blood collected.

The livers were perfused through both the portal vein and hepatic artery using a single inflow tube that was split with a Y-shaped connector. The hepatic vein was connected to the outlet tubing, completing the perfusion loop. After priming the circuit with a peristaltic pump, the pressure required to achieve full perfusion was maintained by a centrifugal pump (Figure 1B). Autologous porcine blood was used as a perfusate, which was maintained at a temperature of 27°C using a heat exchanger (Figure 1B) and oxygenated using a BOC oxygenator (BOC, Oxygen 1-E) with a target Oxygen saturation of 80%. Once the perfusion system was engaged the liver was observed for signs of uniform perfusion with oxygenated blood. Livers were perfused for 30 minutes with pump speeds set at 2500 rotation per minute (RPM) and blood flow maintained at approximately 1000 mL/min (measured by inline sensors on the input stream) in order to overcome the pressure differential between the reservoir in the MRI control room and the organ in the MR scanner. Once stable perfusion of the liver was confirmed, serial MRI images were captured.

**Figure 1.**
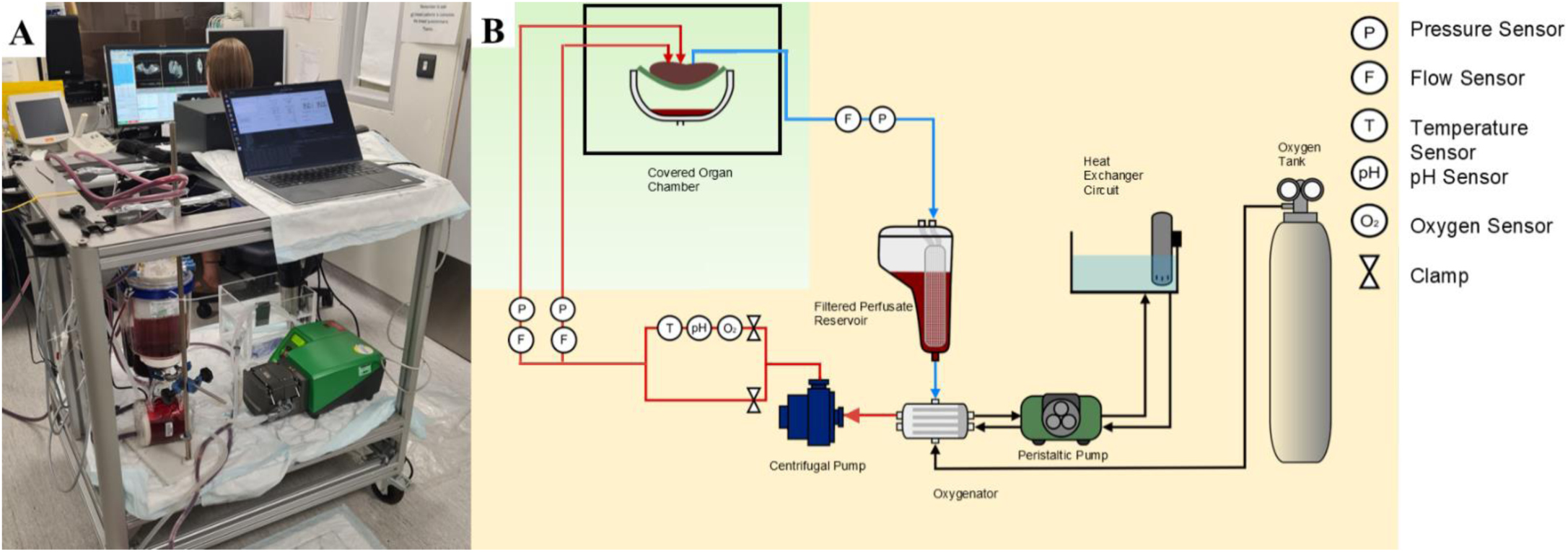
**A**: A photograph of the perfusion rig within the MRI control room (featured: author Dr Caselton). **B**: Schematic of the perfusion circuit: the components in the orange-shaded region are non-MRI-compatible and therefore housed in the EM insulated MRI control room. The light green shaded area demonstrates the non-metallic parts of the system that were within the MRI scanner room (figure adapted from Magbagbeola 2023^10^).

### Magnetic Resonance Imaging (MRI)

MRI was performed using a clinical 3T Philips Ingenia MR scanner (Philips Healthcare, UK). An 18-channel body coil, as used in clinical liver imaging, was positioned on top of the perfused organ chamber for signal acquisition. Respiratory gating was switched off for all sequences. All MRI scans were performed by a trained radiographer with experience in liver imaging, ensuring consistent positioning and optimal image acquisition. A range of clinically relevant liver imaging sequences were employed to acquire structural and functional images.

Anatomical data were acquired with a T_2_-weighted Balanced Turbo Field Echo (BTFE) pulse sequence based on a standard liver imaging protocols (TE = 1.638 ms, TR = 3.2764 ms, flip angle = 30°, echo train length = 274, slice thickness = 5 mm, field of view = 360 x 360 mm).

DWI data were acquired using a spin-echo sequence with b values of 90, 500, 1500 and 2000 s/mm^2^ (TR = 328.19 ms, TE = 75 ms, flip angle = 90°, field-of-view = 348.39 x 348.39 mm, 14 slices with slice thickness = 5 mm), with pulsed diffusion gradients applied separately in three orthogonal directions.

Following the DWI acquisition, DCE-MRI were acquired both prior to and following injection of gadolinium (Gd)-DTPA (Magnevist, Bayer, Germany). 0.05mmoles/kg of Gd-DTPA was injected manually at a rate of 2 mL/s via a cannula placed in the inflow tubing, which connected to the portal vein and hepatic artery, aligning with the dose per kilogram used for in-vivo human studies^30^. DCE-MRI was acquired with a temporal resolution of between 10 and 17 seconds, dependant on field of view.

### MRI Data analysis

Segmentation of blood vessels evident in T2-weighted BTFE images was performed using the ‘magic wand’ region growing tool in Amira 2020.2 (ThermoScientific, UK). Alternating between images acquired in the coronal, sagittal and transverse planes, it was possible to produce a binary segmentation of the vasculature with isotropic voxel dimensions with in-plane resolution (0.83 x 0.83 x 0.83 mm).

A spatial graph of the vessel network was then produced in Amira using the following protocol: (1) a Chamfer distance map was produced from the isotropic segmentation, (2) the Distance-Ordered Thinner module was applied to the distance map with the minimum end length set to two, to avoid erroneous detection of many small branches, (3) the Trace Lines module was used to convert the thinned network into a spatial graph, (4) the spatial graph was smoothed using the Smooth Line Set module (10 iterations, smoothing coefficient = 0.5, attach to data coefficient = 0.25), (5) the Eval of Lines module was used to define vessel radii using a 26-neighbourhood Chamfer distance map for greater accuracy. Graph statistics were exported into Matlab 2023b (Mathworks, USA) for evaluation.

In the case of the DCE-MRI data, gadolinium signal was segmented as above, except in this case the anisotropic voxel dimensions were addressed by resampling the data in the z axis using Amira’s Mitchell algorithm to produce a segmentation with isotropic voxels (between 0.66 x 0.66 x 0.66 mm and 0.83 x 0.83 x 0.83 mm dependant on field of view). A maximum intensity projection through the time dimension of the binary labels (ImageJ) was used to produce a 3D image of the perfused vascular network. From here, the time from the start of the scan to the maximal contrast enhancement was calculated voxel-wise to produce a map of Time To Peak signal intensity (TTP). Concentration enhancement curves were extracted for regions of interest (10 x 10 pixels) that laid entirely within the hepatic vein and portal vein, as close to the inlet/outlet as possible. This was repeated three times for each dataset analysed. The distance moved by the gadolinium signal front was measured manually to estimate flow velocity in the major vessels. Combined with the vessel diameters extracted from the skeletonised data, volumetric flow (*Q*) was calculated according to:

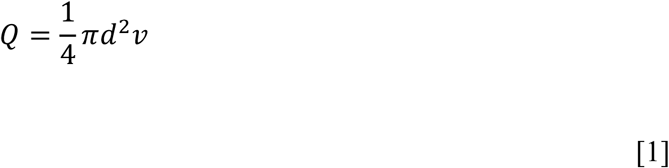

where *d* is vessel diameter and *v* is fluid velocity. The apparent diffusion coefficient (ADC) was calculated pixel-wise from DWI data by fitting the data to a single exponential curve of the form:

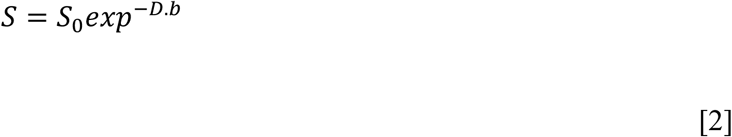

where *S_0_* and *D* are fitted parameters, *S* is the measured signal magnitude. ADC was calculated as the value of *D* averaged across each direction. Mean and standard deviation were calculated for ADC values within the hepatic parenchyma by first subtracting the background and masking the generated ADC maps with the binary vascular segmentation (resampled in ImageJ to aligned with DWI data) to exclude values measured within the vessels. ImageJ’s histogram function was then used to plot the distribution of pixelwise ADC values and find the mean and standard deviation.

### Histological evaluation

Histological evaluation was performed on biopsies taken from the livers stored using SCS (control) and the perfused and imaged livers to assess whether preservation-related or perfusion-related injury had occurred. Six tru-cut biopsies were taken at random sites within each sample, three prior to perfusion or storage (baseline), and a further three following the 30 minute perfusion or storage period. Immediately after collection, liver biopsy tissues were immersed in 10% neutral-buffered formalin for a minimum of 24 hours to ensure fixation. Following this, they were processed using an automated tissue processor, which involved sequential dehydration in graded ethanol solutions, clearing in xylene, and infiltration with molten paraffin wax. Once processed, the tissues were embedded in paraffin blocks, ensuring proper orientation. These blocks were then sectioned at a thickness of 5 μm using a rotary microtome, with sections floated on a water bath at 42°C and retrieved on glass slides. For staining, slides were first deparaffinized in xylene, then rehydrated through a graded ethanol series, and stained with hematoxylin for 5 minutes. After rinsing, they were counterstained with eosin for 3 minutes. Following staining, the slides underwent dehydration through graded ethanol, cleared in xylene, and mounted using Permount Mounting Medium (Fisher Chemical, UK).

Histology samples were evaluated according to a bespoke scoring system, which was created to quantitatively assess key histopathological features linked to reperfusion injury and overall liver health. The parameters for assessment (neutrophil infiltration, hepatocyte necrosis, sinusoidal congestion, and oedema) were chosen to provide an integrated perspective on tissue changes. A score between zero and three was awarded in each category according to the most severe observation within the examined field (Table 1). The total pathology score, ranging from zero to 12, was derived by summing the scores for individual features.

**Table 1:**
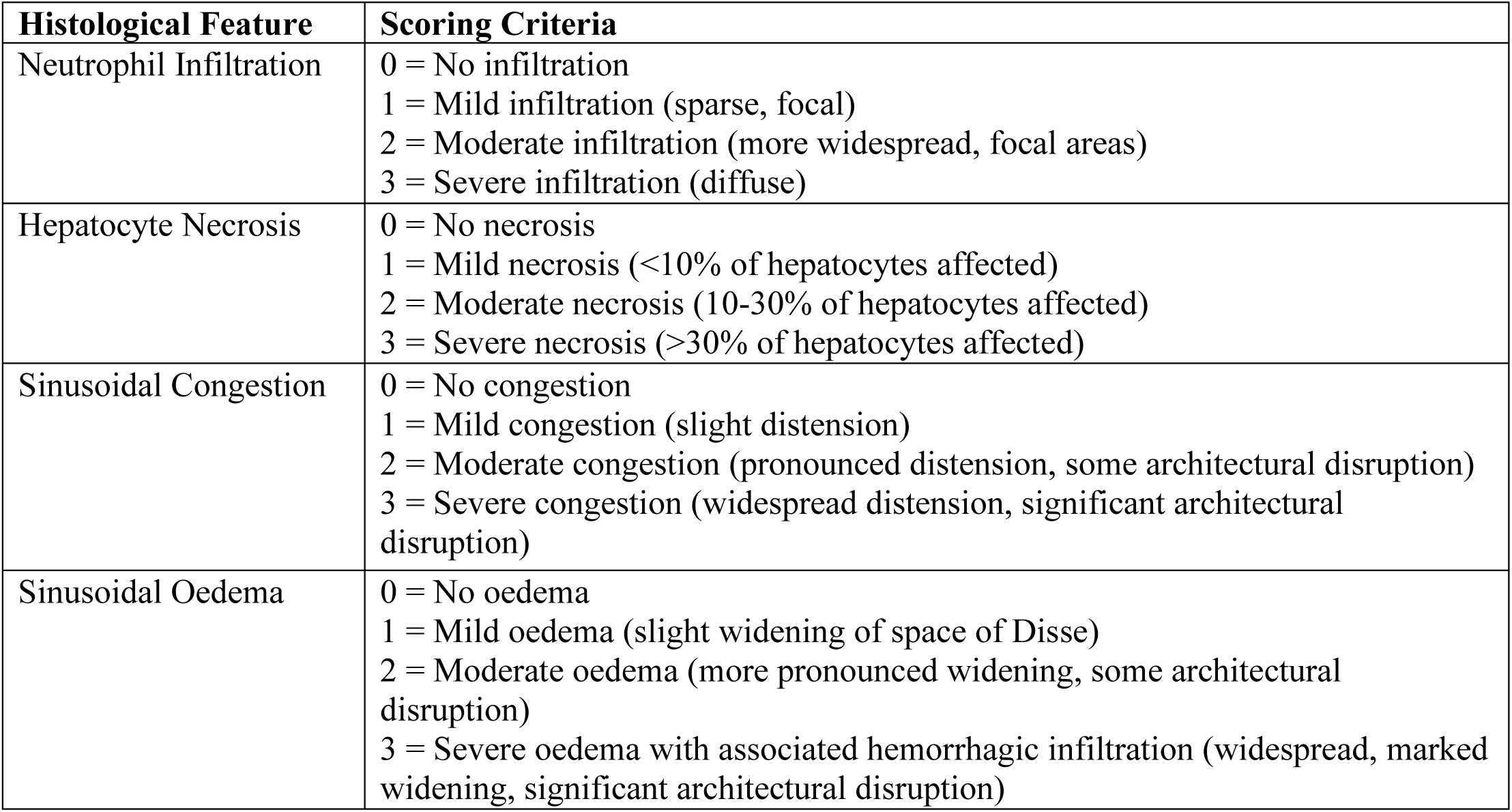
Custom Scoring System for Histological Assessment of Acute Liver Pathology.

| Histological Feature | Scoring Criteria |
| --- | --- |
| Neutrophil Infiltration | 0 = No infiltration<br>1 = Mild infiltration (sparse, focal)<br>2 = Moderate infiltration (more widespread, focal areas)<br>3 = Severe infiltration (diffuse) |
| Hepatocyte Necrosis | 0 = No necrosis<br>1 = Mild necrosis (<10% of hepatocytes affected)<br>2 = Moderate necrosis (10-30% of hepatocytes affected)<br>3 = Severe necrosis (>30% of hepatocytes affected) |
| Sinusoidal Congestion | 0 = No congestion<br>1 = Mild congestion (slight distension)<br>2 = Moderate congestion (pronounced distension, some architectural disruption)<br>3 = Severe congestion (widespread distension, significant architectural disruption) |
| Sinusoidal Oedema | 0 = No oedema<br>1 = Mild oedema (slight widening of space of Disse)<br>2 = Moderate oedema (more pronounced widening, some architectural disruption)<br>3 = Severe oedema with associated hemorrhagic infiltration (widespread, marked widening, significant architectural disruption) |

## Results

### Organ retrieval and preservation periods

A total of 14 porcine livers were harvested and used in the study. Seven were subjected to MRI-compatible sub-normothermic machine perfusion, with the other seven serving as temporaneous controls. Three of the 7 livers harvested and perfused were rejected due to parenchymal tears sustained in the abattoir which could not be adequately repaired.

### Confirmation of successful perfusion

Satisfactory establishment of the perfusion circuit in the machine-perfused livers was confirmed by visual inspection and supported by flow monitoring using inbuilt flow sensors. Inspection assessed signs of even colouration and absence of haemorrhage, before the organ was placed in the MR scanner (Figure 2A-C). Flow sensors enabled continuous monitoring of perfusion whilst the organ was within the MRI scanner: a decrease in measured outflow rate relative to inflow rate indicated a potential leak requiring repair (Figure 2D). Four out of the seven perfused livers achieved successful sustained perfusion, defined by the establishment of a perfusion circuit followed by a 30-minute continuous perfusion. The perfusate was observed in the reservoir for signs of clotting, which would negatively impact perfusion, however it was found that the heparinised perfusate prevented autologous blood from clotting during the perfusion period.

**Figure 2.**
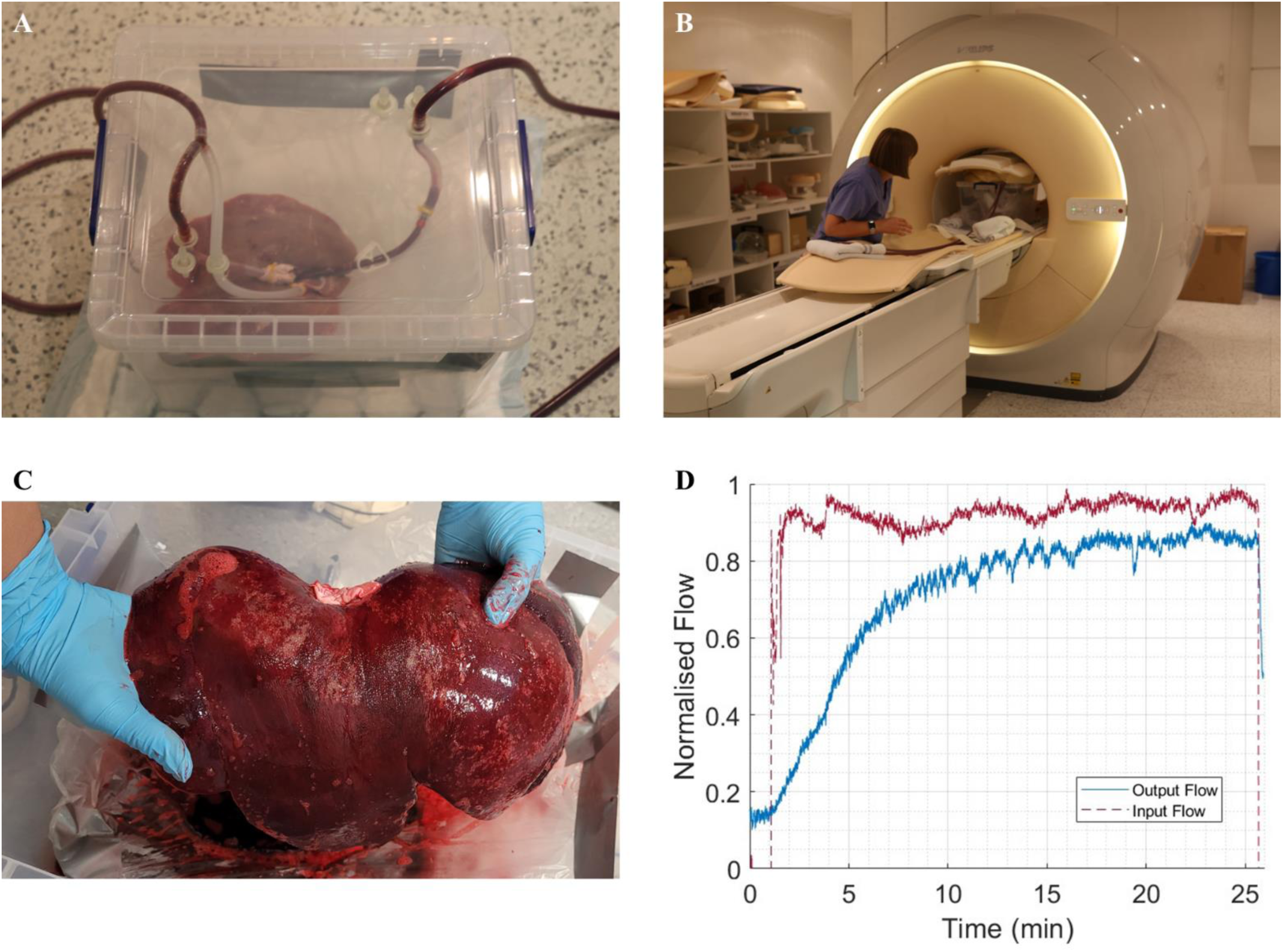
**A**: A photograph of a liver being primed for perfusion within the water-tight organ housing. **B**: The primed organ is photographed inside the clinical MR scanner. **C**: A liver is shown following successful perfusion. The even, dark colouration was used as indicator of successful perfusion alongside sensor outputs. **D**: Representative flow sensor output from a successful 25-minute perfusion period within the MR scanner. The input flow rate (red) is maintained at a steady level, and the output flowrate is monitored to ensure the perfusion circuit is not losing significant perfusate, which would be indicative of a leak. MRI scans were acquired once outflow had stabilised.

### Histology

Histological analysis of ischemia and reperfusion injury was performed on the perfused livers (n=4) and compared with those stored in Static Cold Storage (SCS) (n=4) at baseline and following 30 minutes of perfusion. A custom scoring system was used to assess tissue damage, and scores were averaged across the four experiments (Table 2). At baseline, there was no statistically significant difference between the groups (p = 0.704, independent t test). However, after 30 minutes of perfusion or SCS, a significant difference in tissue preservation was observed, with perfused livers displaying less tissue degradation (mean composite score = 8.4) compared to the SCS group (mean composite score = 11.9, p = 0.024). Data normality and homogeneity of variances were confirmed using Shapiro-Wilk test (P > 0.05) and Levene’s test (0 > 0.05) respectively.

**Table 2:** Mean histological scores for liver samples at different time points. The scores are calculated on a scale of 0–3 for each category, where 0 represents no change or damage, and 3 reflects severe change or damage. The composite score is obtained by summing the four categories.

| Group | Time Point | Mean Neutrophil Infiltration | Mean Hepatocyte Necrosis | Mean Sinusoidal Congestion | Mean Sinusoidal Oedema | Mean Composite Score |
| --- | --- | --- | --- | --- | --- | --- |
| Perfused | Baseline | 1.0 | 1.3 | 2.1 | 1.9 | 6.3 |
| SCS | Baseline | 1.0 | 1.6 | 2.0 | 2.0 | 6.6 |
| Perfused | 30 min | 1.1 | 2.2 | 2.1 | 3.0 | 8.4 |
| SCS | 30 min | 2.9 | 3.0 | 3.0 | 3.0 | 11.9 |

Representative histological images demonstrate the well-maintained cellular architecture seen in the perfused group (Figure 3A and C), compared to the SCS livers, where there was marked oedema and a loss of hepatic cellular architecture (figure 3B and D).

**Figure 3.**
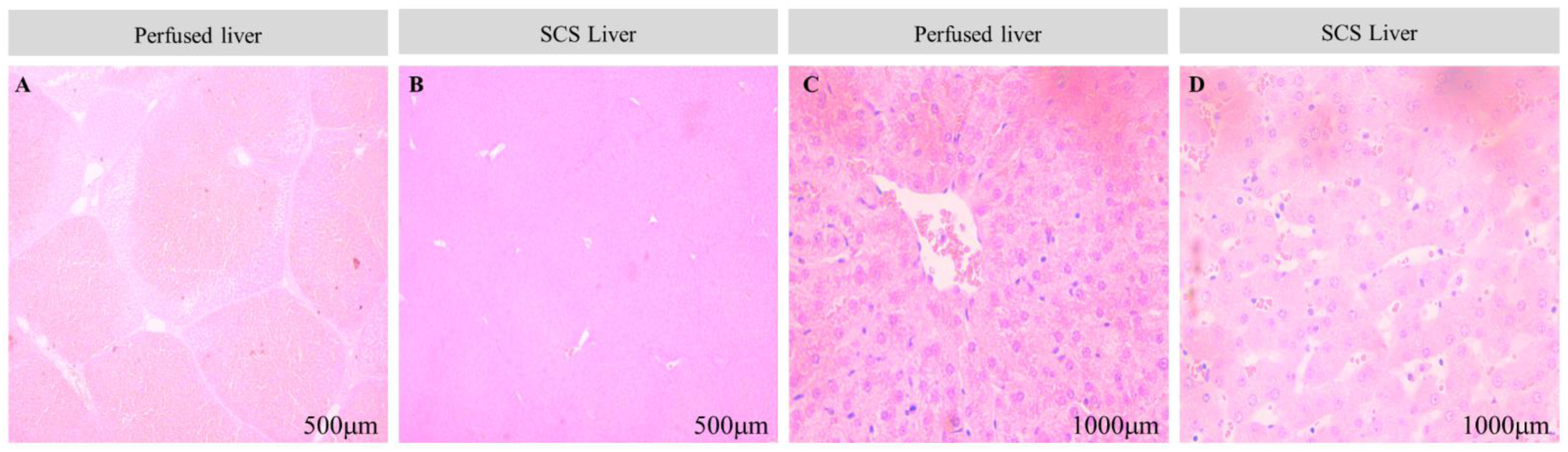
**A**: Perfused liver at 30 minutes following start of perfusion demonstrating clear well-demarcated hepatic acini. **B:** SCS liver at 30 minutes following start of perfusion. Compared to the perfused liver the hepatic acinar structure is less well-defined. **C:** Higher magnification of the perfused liver demonstrates intact hepatocyte cellular architecture, compared to the loss of cellular organisation seen in the SCS control, **D**.

### T2-weighted imaging of liver morphology

T2-weighted images of the perfused livers showed clear anatomical landmarks (Figure 4A) demonstrating that the perfusion chamber did not impede imaging. In some samples, the inflow and outflow tubes were also visible, providing confirmation of correct positioning within the portal vein, hepatic vein and hepatic artery (Figure 4B). The portal vein and hepatic vein are clearly identifiable, allowing the venous networks to be segmented (Figure 4C). T2 weighted images were acquired in the coronal, sagittal and transverse planes sequentially, and then resampled to produce isotropic voxels, facilitating registration of the orthogonal views. This allowed concurrent segmentation of vessels in the three planes, capitalising on the high in-plane resolution to produce isotropic labels of the venous networks down to 0.45 mm in radius (Figure 4D-E). However, intra-hepatic branches of the hepatic artery were not able to be traced.

**Figure 4.**
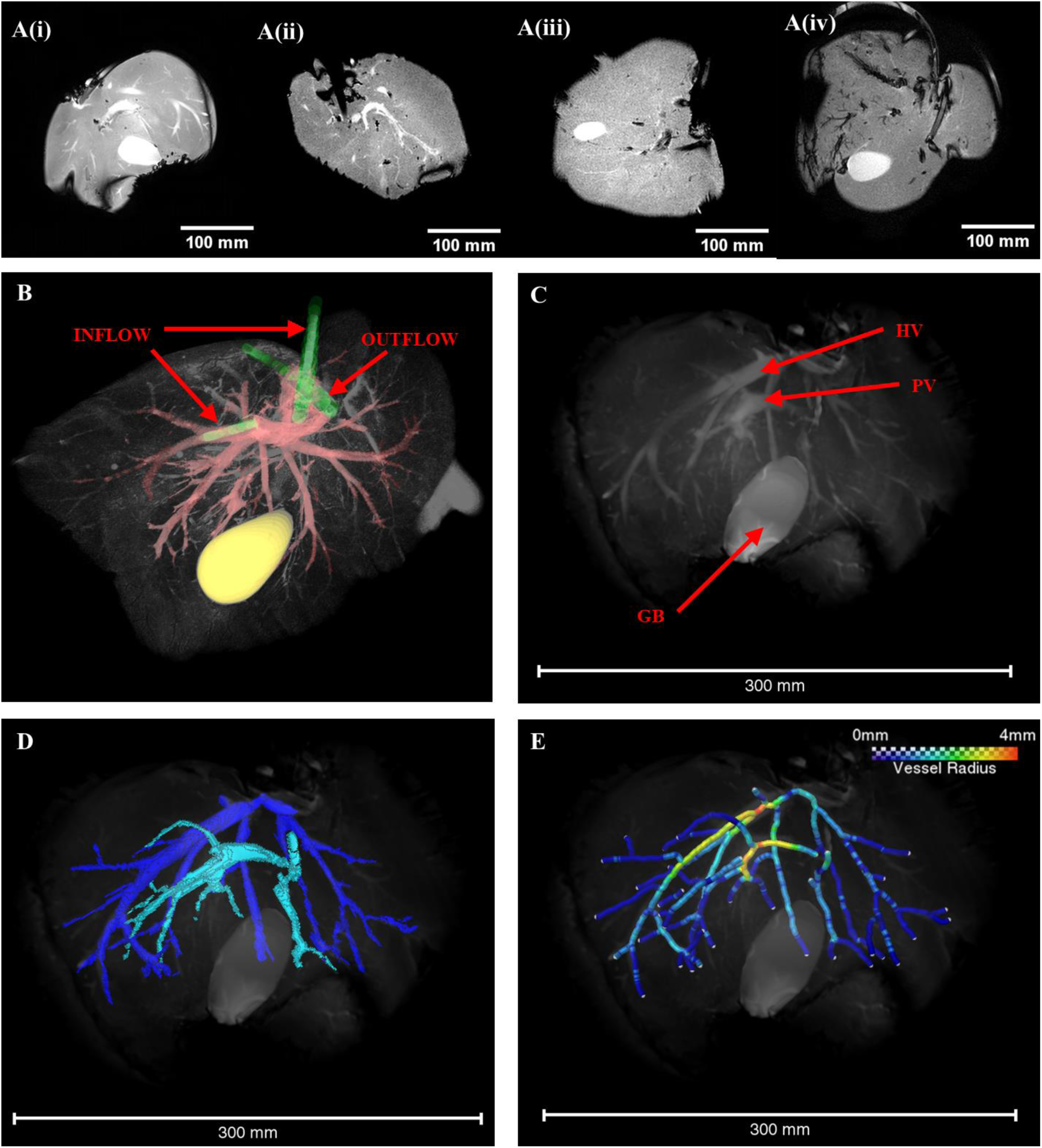
**Ai-iv**: The central 2D slice from BTFE T2 weight coronal MRI images are shown for four successfully perfused livers. **B**: A 3D rendered T2 weighted image of a different liver is shown with the inflow and outflow tubing highlighted in green. Tubing can be seen entering the portal vein and hepatic artery (inflow) and the hepatic vein (outflow). Vessels are highlighted in red in this image, and the gall bladder in yellow. **C**: A 3D rendering of a BTFE T2 weighted MRI image, resampled in the z axis to produce isotropic voxels. Key anatomical landmarks are indicated with red arrows: the gall bladder (GB), hepatic vein (HV) and portal vein (PV). **C**: Labelled vasculature was segmented from three orthogonal BTFE MRI images. The high in-plane resolution of each orthogonal image aided in construction of the two separate venous vessel trees: the branches of the hepatic vein are labelled in dark blue and the portal vein in light blue. **D:** A skeletonised vessel network was extracted from the segmentation shown in (B). Segment colour indicates the vessel radius, with the largest vessels in this sample having a radius of 4 mm. The smallest identifiable vessels had a radius of 0.45 mm.

From the labelled images, it was possible to extract a variety of metrics describing the vascular morphology of the four livers studied. Comparing the skeletonised networks, it was found that the total vascular volume and total vessel length varied between samples (Figure 5A), with one liver displaying an especially wide branch of the hepatic vein (radius 8.0mm) leading to a high total vessel volume. All four livers had an approximately even ratio of vessel branching points to terminal nodes within their networks (45-53% terminal nodes, Figure 5B), and the majority of branching points represented bifurcations (mean coordination numbers ranging from 3.02 to 3.23 across all samples). The median length of vessel segments (defined as the curved length of a vessel between two nodes) ranged between 4.2 and 14.9 mm for the four samples, with liver sample 4 displaying on average longer vessel segments and fewer nodes (Figure 5C). The median vessel radius ranged between 0.8 and 1.3 mm (Figure 5D). Vessel tortuosity was calculated from the curved segment length divided by the straight length, and in all four livers the median tortuosity was between 1.04 and 1.05.

**Figure 5.**
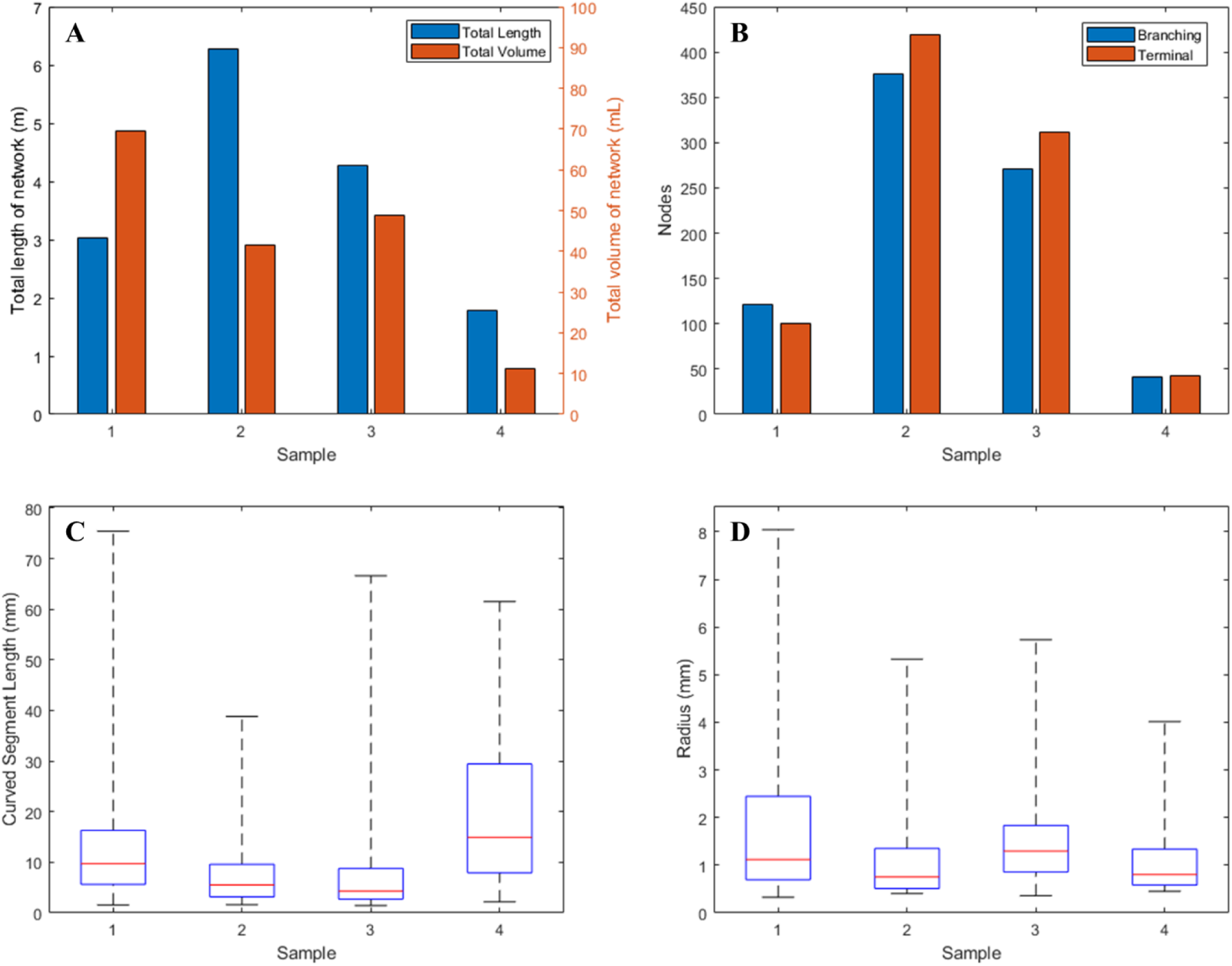
**A**: Total vessel length and volume extracted from four livers (sample 1-4). Variation is seen between the livers, particularly in terms of total vessel volume. **B**: The number of branching and terminal nodes in each liver network is shown. While the number of nodes varies, the ratio is similar across all four livers. **C**: The distribution of curved vessel segment lengths is shown as a boxchart (minimum and maximum values denoted by black whiskers, interquartile range shown as a blue box, median value shown as a red line). **D**: The distribution of vessel radii is shown for each liver sample. The minimum radius size was limited by the imaging resolution, while the maximum vessel radius was between 4 mm and 8 mm. In all cases the largest vessel radius was found in the hepatic vein.

### Diffusion Weighted Imaging

DWI data was acquired with b values of 90, 500, 1500, and 2000 s/mm^2^ in three of the four perfused livers, and analysed to produce ADC maps (Figure 6). The mean ADC within the parenchyma was found to be 0.55×10^-3^ s/mm^2^ (standard deviation 0.36×10^-3^ s/mm^2^). These values on perfused porcine livers fall at the lower end of the range reported for human liver parenchyma in the literature (0.69-2.28×10^-3^ s/mm^2^)^31^.

**Figure 6.**
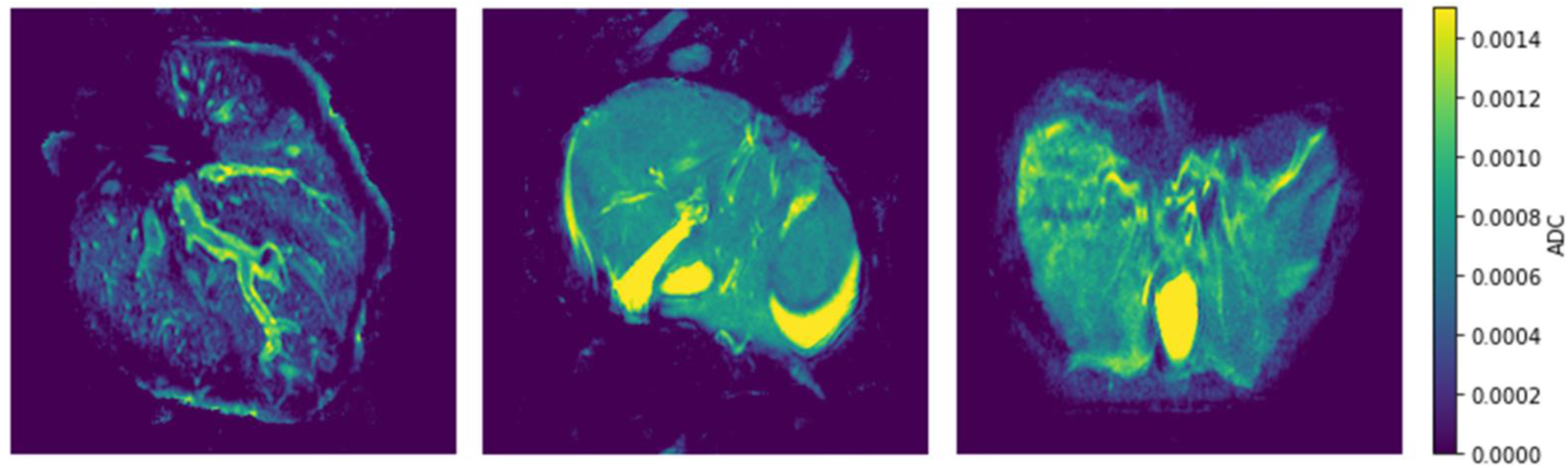
These representative 2D sections show the ADC maps calculated for three perfused livers. Higher apparent diffusion (yellow) is seen in the gall bladder and vasculature, compared to the hepatic tissue (blue). (units = s/mm^2^)

### Dynamic Contrast Enhanced MRI

DCE images were acquired following Gadolinium administration into the perfusion circuit at the inlet tube. The contrast agent could be seen moving from the portal vein, through the circulatory network, and into the hepatic vein over the course of 5 minutes (Figure 7). This confirms that the perfusion circuit successfully set up a circular flow.

**Figure 7.**
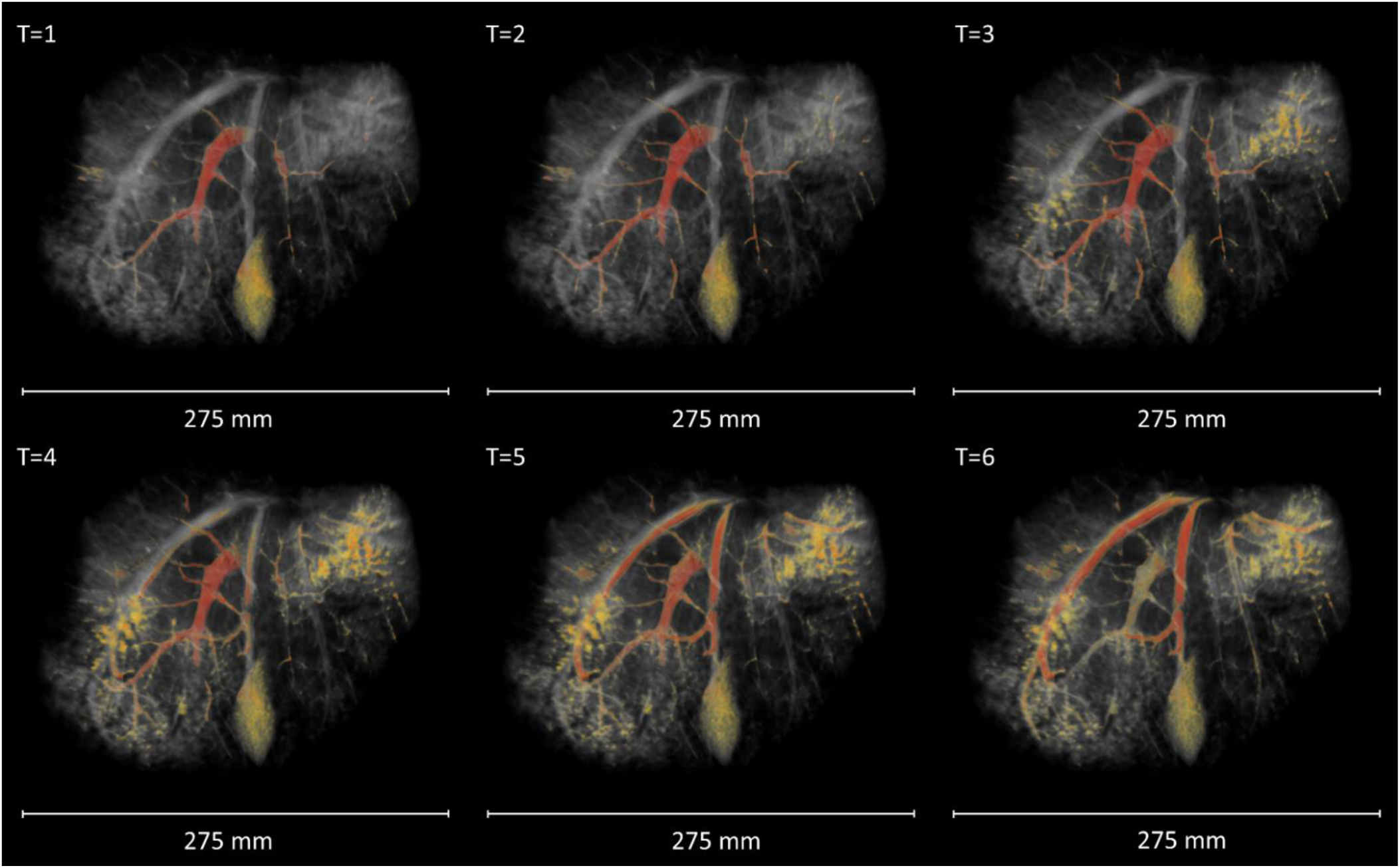
Perfusion of contrast agent through the vasculature, imaged using DCE-MRI. The gadolinium signal at each time point is highlighted in red. The signal enhancement is first confined to the portal vein (T1-2), then spreads to the smaller vessels and extra-vascular tissue (T3-4). Finally, the branches of the hepatic vein are enhanced (T5-6). This indicated that the perfusion circuit is working as intended.

Time To Peak signal (TTP), a semi-quantitative measure of flow dynamics, was calculated voxel-wise (Figure 8A-B). Concentration enhancement curves were calculated at the point of widest radius within the portal vein and hepatic vein (Figure 8C-D). Variation was seen between livers, with one liver showing concurrent enhancement of the hepatic vein and portal vein (time to maximum enhancement of 86.9 and 97.7 seconds respectively). The remaining livers demonstrated early enhancement in the portal vein (times ranging from 27.5 to 97.7 seconds post-injection) and delayed hepatic vein enhancement (between 97.9 and 179.1 seconds post-injection). The time delay between peak signal in the portal and hepatic veins varied between 69.9 and 81.4 seconds in these livers.

**Figure 8.**
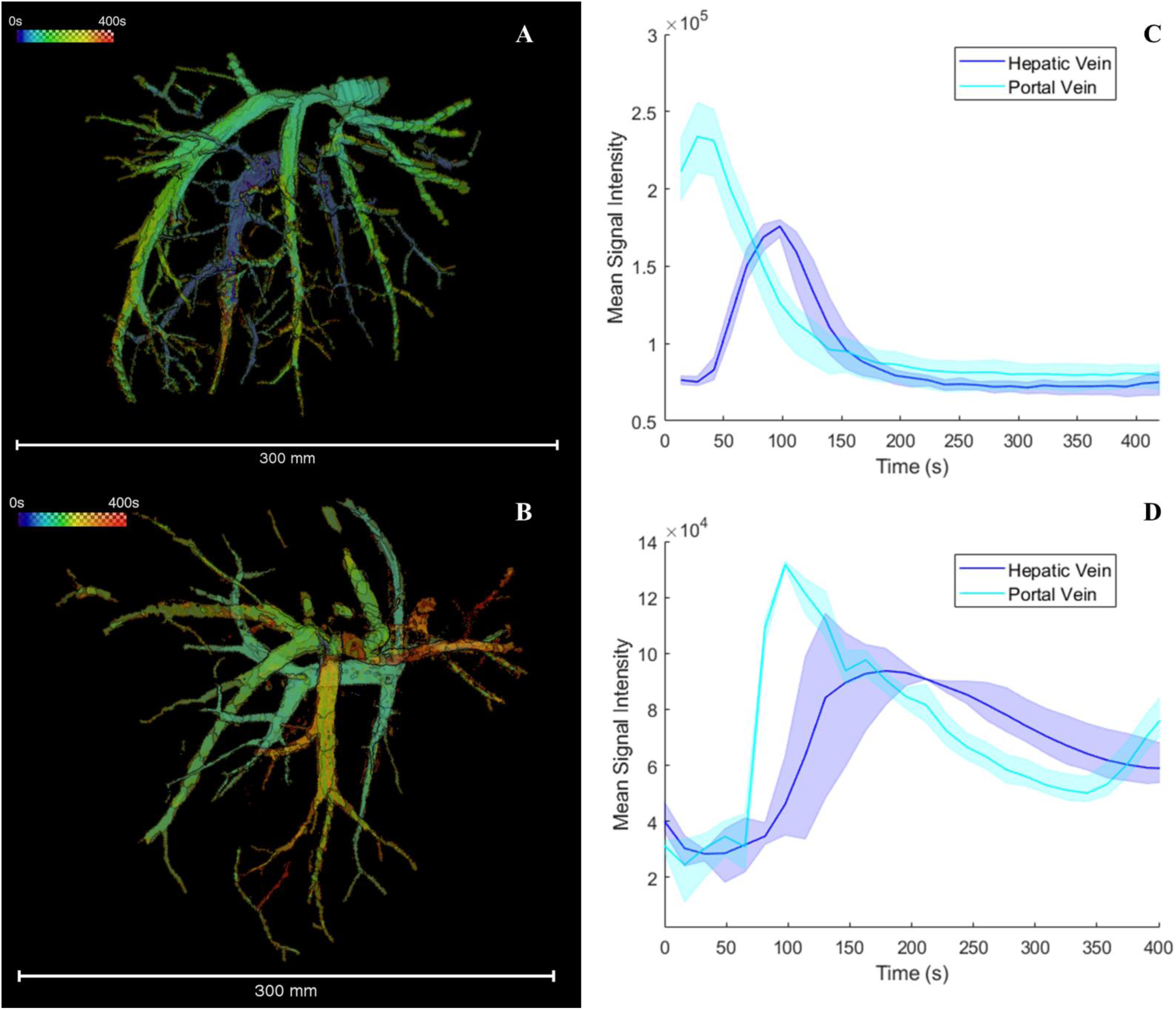
**A-B**: The vascular network of two perfused livers shown colour-coded according to the Time To Peak signal (TTP), calculated voxel-wise. **C-D**: Contrast enhancement curves are shown for two regions of interest, taken from the portal vein (dark blue) and hepatic vein (light blue) respectively. The shaded region denotes the maximum and minimum contrast signal recorded at each timepoint from the three repeat readings, while the solid line shows the mean signal. As expected, the gadolinium signal quickly reaches the inlet vessel, while the hepatic vein shows a delayed contrast enhancement.

Additional flow analysis was conducted on one liver sample to calculate the flow rate within the branches of the hepatic vein and the portal vein (Figure 9). The flow speed, calculated as the distance moved by the gadolinium bolus divided by time, was similar in the portal and hepatic veins (31.25 +/- 3.55 cm/min and 28.18 +/- 8.37 cm/min respectively). However, the flow rate was higher in the hepatic vein: 28.39 +/- 6.54 mL/min compared to 4.86 +/- 0.86 mL/min in the portal vein.

**Figure 9.**
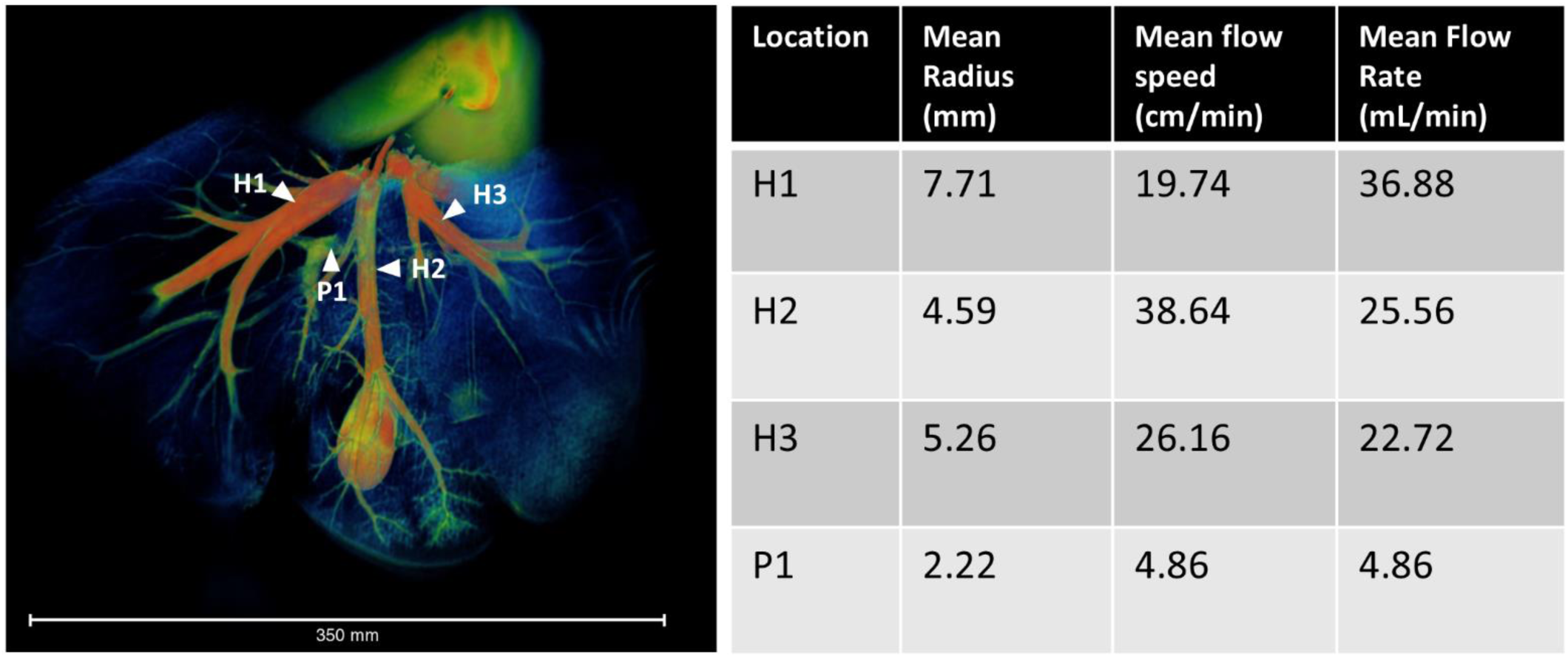
**A**: A 3D rendering of a perfused liver. Numbered white arrows indicate locations at which flow velocity was measured within the hepatic vein (H1-3 for the three major branches) and portal vein (P1). **B**: A table showing mean vessel radius, flow speed and flow rate at each measurement location. A higher flow rate is seen in the hepatic veins compared to the portal vein due to the smaller vessel radius measured in the portal vein.

## Discussion

This if the first study to achieve quantitative flow characteristics using MRI of a healthy ex vivo liver facilitated by an ex vivo organ perfusion system developed for experimental use. The information achieved would suggest a huge potential in assessing the vascular changes associated with chronic liver disease, liver transplantation, liver cancer and the effect and optimisation of agents used in their treatment.

### Investigation of vascular morphology

Hepatic blood vessel networks were segmented at high resolution in three orthogonal planes to achieve isotropic resolution, allowing for more accurate analysis of vascular morphology. Morphological analysis highlighted the variability of total vascular volume and vessel length between liver samples. Vessel radii and segment length were measured, and parameters such as vessel tortuosity were extracted. These metrics are important for future application of perfused organs as a model of disease, as changes in vascular morphology can be indicative of disease progression and treatment response. Vessel tortuosity, for example, is a hallmark of many cancer types^32^ among other conditions^33^, while liver pathologies such as cirrhosis are known to cause changes in vessel diameter^34^. Furthermore, the extracted vessel network graphs can be used in image-based computer modelling of flow and solute delivery, enabling research into the effects of vascular morphology in disease^35,36^. Nonetheless, there are limitations to this methodology: most notably, it was not possible to trace hepatic artery branches in these images, most likely due to their small diameter in comparison to portal and hepatic vein branches. Administration of a contrast agent to the hepatic artery inflow only could be used to highlight these vessels.

### Measurements of microcirculation and flow

DWI data was acquired and used to calculate ADC values for three of the four successfully perfused livers, with one liver being excluded from the analysis due to poor image quality. The measured ADC values (mean 0.55×10^-3^ s/mm^2^, standard deviation 0.36×10^-3^ s/mm^2^) were within the range of values for human parenchyma reported in literature^31^. This result suggests that machine perfused porcine livers have similar MRI characteristics to the human liver and could be used to study vascular changes in human disease. Notably, the use of machine-perfused organs here avoided the need for the respiration gating or repeated breath-holds that would be required in live or anaesthetised subjects. As these are ex vivo perfused livers, it was not possible to compare the ADC values against measurements in another organ, for example the spleen, as recommended for clinical DWI^37,38^. Moreover, measured ADC can be dependent on scan parameters, so quantitative comparisons to literature are limited^39,40^.

DCE-MRI was successfully performed in the perfused livers with contrast administered via a cannula in the perfusion inflow tube. Image acquisition over a period of approximately 5 minutes allowed visualisation of contrast agent progression through the hepatic vasculature. The time to peak contrast enhancement and concentration enhancement curves for regions of interest within the portal and hepatic veins showed a directional flow from the portal vein to the hepatic artery, except for one sample which saw simultaneous contrast enhancement within both portal and hepatic veins which was attributed to a parenchymal injury within the liver resulting in vascular shunting.

Measuring the distance advanced by the gadolinium bolus front allowed for the calculation of flow rate within the major vessels, with a mean flow rate of 28.39 +/- 6.54 mL/min in the hepatic vein, and 4.86 +/- 0.86 mL/min in the portal vein. To reduce subjectivity, a semi-automated intensity-based algorithm was used in Amira to detect the front, but which still required substantial manual input.

Whilst machine parameters will influence both DWI and DCE-MRI results, these results demonstrate that these two widely used clinical measurements are feasible within this system, and exploring the effects of varying perfusion parameters, such as flow rate and pressure, on MRI results could lead to new roles in clinical imaging.

### Organ collection and perfusion within the scanner

The advantages and limitations of acquiring organs from an abattoir are discussed in detail in a previous work^11^. In brief, the decision to make use of fresh porcine livers collected from an abattoir for this study was based on the advantages of low cost, high availability and the ethical consideration of using organs from animals already scheduled for slaughter. However, an important challenge is to avoid damage to the liver during en bloc removal of the abdominal organs in the abattoir. Three of the seven livers collected for this study were damaged and proved to be unsuitable for perfusion. When perfusing the livers within the MRI scanner, an increase in failure of the perfusion circuit (specifically leaks) was noted when compared to previous studies using the same custom perfusion rig^10,11^. This was likely due to the increased pressures required to circulate the perfusate through the extended length of tubing between the consol room and MRI scanner.

### Organ Viability

Previous research within our group has demonstrated that the perfusion system can maintain whole porcine livers in a viable state for up to 3 hours^10^. Histological analysis in this study revealed intact hepatocyte cellular architecture and lack of oedema compared to SCS controls, indicating minimal damage to the liver microstructure during our experiments. For longer-duration experiments in future studies, this could be complemented by measurement of metabolic data^22,41^.

### Applicability to human and pathological organs

The experiments were performed in healthy porcine livers, which served as a useful model organ. The results provide a promising proof-of-concept to guide research application in human organs such as livers deemed unsuitable for clinical transplant or partial human liver resections performed to remove primary or secondary liver cancers. Given the inherent anatomical and physiological differences between porcine and human livers, we expect that further optimisation of the methodology will be required.

## Conclusion

In summary, this work demonstrates the use of machine-perfused viable whole liver combined with MRI as a research platform for studying flow in intact organs. We have demonstrated the potential of structural and functional MRI sequences for quantitatively investigating vasculature morphology and perfusion in machine-perfused organs and, by employing imaging sequences regularly used in a clinical setting for assessment of liver disease, we extracted physiologically relevant parameters relating to microcirculation and flow. While this study used healthy porcine livers, further work will extrapolate these methods to human and diseased tissues. Using these methods, it will be possible to investigate pathological physiology in human tissues under a controlled environment, as well as evaluate medical interventions relating to blood flow and analyse the distribution and pharmacokinetic properties of drugs.

## Abbreviations

MRI: Magnetic Resonance Imaging
DCE: Dynamic Contrast Enhancement
DWI: Diffusion Weighted Imaging
ADC: Apparent Diffusion Coefficient
BTFE: Balanced Turbo Field Echo

